# Intelligent Design of *Escherichia coli* Terminators

**DOI:** 10.1101/2025.03.04.641434

**Authors:** Jie Li, Lin-Feng Wu, Kai Liu, Bin-Guang Ma

**Author notes:** Corresponding author., E-mail address (Bin-Guang Ma). Equal contribution (Jie Li, Lin-Feng Wu).

## Abstract

Terminators are specific nucleotide sequences located at the 3’ end of a gene and contain transcription termination information. As a fundamental genetic regulatory element, terminators play a crucial role in the design of gene circuits. Accurately characterizing terminator strength is essential for improving the precision of gene circuit designs. Experimental characterization of terminator strength is time-consuming and labor-intensive; therefore, there is a need to develop computational tools capable of accurately predicting terminator strength. Current prediction methods do not fully consider sequence or thermodynamic information related to terminators, lacking robust models for accurate prediction. Meanwhile, deep generative models have demonstrated tremendous potential in the design of biological sequences and are expected to be applied to terminator sequence design. This study focuses on intelligent design of *Escherichia coli* terminators and primarily conducts the following research: (1) To construct an intrinsic terminator strength prediction model for *E. coli*, this study extracts sequence features and thermodynamic features from *E. coli* intrinsic terminators. Machine learning models based on the selected features achieved a prediction performance of *R*^2^ = 0.72. (2) This study employs a generative adversarial network (GAN) to learn from intrinsic terminator sequence training data and generate terminator sequences. Evaluation reveals that the generated terminators exhibit similar data distributions to intrinsic terminators, demonstrating the reliability of GAN-generated terminator sequences. (3) This study uses the constructed terminator strength prediction model to screen for strong terminators from the generated set. Experimental verification shows that among the 18 selected terminators, 72% exhibit termination efficiencies greater than 90%, confirming the reliability of the intelligent design approach for *E. coli* terminators. In sum, this study constructs a terminator strength prediction model and a terminator generation model for *E. coli*, providing model support for terminator design in gene circuits. This enhances the modularity of biological component design and promotes the development of synthetic biology.

## 1. Introduction

A terminator is a specific nucleotide sequence located downstream of a gene that provides termination information for the transcription of that gene [1]. Based on the forms of transcription termination in prokaryotes, terminators can be classified into rho-dependent terminators and intrinsic terminators [2]. Rho-dependent terminators achieve transcription termination by relying on the interaction between the rho factor and RNA polymerase to terminate transcription upstream of the gene [3]. Intrinsic terminators primarily consist of an RNA secondary structure, which includes a 5’ poly-A tract, a hairpin structure, and a 3’ poly-U tract. The hairpin structure is composed of GC-rich stem regions and loop regions [4]. To date, studies have characterized terminator strength in prokaryotes, and researchers have constructed terminator strength prediction models based on these characterization data [5-7]. However, current methods for predicting terminator strength do not fully utilize the sequence and thermodynamic information of terminators, particularly failing to account for the individual sequence features of each region of the RNA secondary structure. As a result, accurate prediction models for terminator strength remain lacking.

Repeating the use of terminator sequences within the same genetic circuit may lead to homologous recombination events, resulting in genetic instability. Additionally, traditional design methods significantly increase the number of potential sequences that need to be screened as sequence length increases [8]. For example, for a promoter sequence of 50 bp in length, there are over 10^30^ potential sequence combinations, which far exceeds experimental screening capacity. At present, generative adversarial networks (GANs) have been widely applied to the design of biological sequences [9-15] and have demonstrated remarkable capabilities. By exploring the latent space of existing biological sequences, GANs learn the data distribution characteristics of these sequences and generate new biological sequences with similar distributions, thereby enabling the selection of sequences that meet specific criteria [16, 17].

In this study, we constructed a terminator strength prediction model based on sequence features, thermodynamic features, and RNA secondary structure sequence features of *E. coli* intrinsic terminators. Since strong transcription termination is often required in genetic circuit design to avoid interference between transcription units, particularly when high promoter activity or compact arrangement of circuit components is needed [18], we employed deep generative models (GANs) to generate terminator sequences. We then used the *E. coli* terminator strength prediction model to screen for strong terminators from the generated sequences and experimentally measured their termination efficiencies to validate the reliability of both the prediction model and the generation model.

## 2. Materials and methods

### 2.1 Construction of terminator strength prediction model

#### 2.1.1 Data Source

The experimental dataset on *E. coli* terminator strength used in this study was constructed by Chen et al. [6]. This dataset contains 607 *E. coli* terminators and their corresponding termination strength data. To validate the robustness and generalization ability of the terminator strength prediction model developed in this study, the dataset was randomly divided into a training set and a test set at a 4:1 ratio.

#### 2.1.2 Feature extraction

##### Sequence features

The sequence features used in this study include *k*-mer, C*K*SNAP (Composition of *K*-spaced Nucleic Acid Pairs) [19], and Pseudo *K*-tupler Nucleotide Composition (Pse*K*NC) [20]. The *k*-mer feature represents the frequency of adjacent *k*-length nucleotide sequences in a given nucleotide sequence. Its calculation formula is:

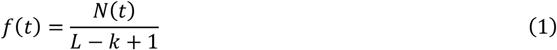

Here, *t* denotes a specific *k*-tuple nucleotide sequence (with 4^*k*^ possible combinations), *N*(*t*) represents the occurrence count of a specific *k*-length substring in the sequence, and *L* is the total sequence length [21]. C*K*SNAP characterizes the frequency of dinucleotide pairs separated by *K* nucleotides, where *K* is an integer ranging from 0 to 5 (default maximum *K* = 5) [19]. When *K* = 0, C*K*SNAP corresponds to the 2-mer feature in *k*-mer. The C*K*SNAP formula is:

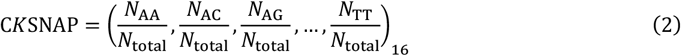

For a nucleotide sequence of length *L, N*_total_ equals *L*−1, *L*−2, *L*−3, *L*−4, *L*−5, and *L*−6 when *K* = 0, 1, 2, 3, 4, 5, respectively.

*PseKNC* is a sequence feature extraction method that incorporates both short-range and long-range sequence information [20]. By integrating physicochemical properties with *k*-tuple nucleotide composition, this method effectively captures long-range dependencies within the sequence.

##### Thermodynamic features

The thermodynamic features of terminators in this study are characterized based on the RNA secondary structures formed during transcription. The thermodynamic features used are listed in **Table 1**.

**Table 1.** Description of thermodynamics related features.

| Thermodynamic Feature | Description |
| --- | --- |
| $\Delta G_T$ | Minimum free energy of the terminator sequence |
| $\Delta G_U$ | Free energy of the U-tract structure in the terminator sequence |
| $\Delta G_L$ | Free energy of the loop structure in the RNA hairpin of the terminator |
| $\Delta G_H$ | Free energy of the RNA hairpin structure formed by the terminator sequence |
| $\Delta G_H/\text{Length\_Hairpin}$ | Free energy of the hairpin structure divided by its sequence length |
| $\Delta G_H/\text{Length\_Stem}$ | Free energy of the hairpin structure divided by the stem sequence length |
| $\Delta G_B$ | Free energy of the bottom three base pairs in the stem of the hairpin structure |

The $\Delta G_T$ , $\Delta G_L$ , $\Delta G_H$ and $\Delta G_B$ values for terminator sequences were calculated using the Vienna RNA package toolbox [22].

The free energy of the U-tract structure, Δ*G*_*U*_, represents the binding energy between the U-tract of the terminator and the template DNA. The weak base pairing of this structure with the template DNA facilitates the dissociation of the terminator from the DNA [6]. The formula for Δ*G*_*U*_ is as follows:

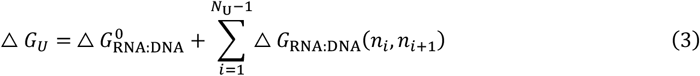

In this formula, when *N*_U_ = 8, it indicates the calculation of the binding energy between the first 8 bases of the U-tract and the template DNA. Here, 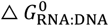 denotes the initial free energy of the RNA:DNA hybrid strand, andΔ *G*_RNA:DNA_(*n*_*i*_, *n*_*i*+1_) represents the free energy of the RNA:DNA hybrid strand extending from position *i* to *i*+1 in the U-tract.

##### Sequence features of terminator RNA secondary structures

This study extracts sequence features based on the RNA secondary structures formed by terminator sequences during transcription. These features include the following:

1. U-tract structure’s *U*_score_. *U*_score_ characterizes the distribution of U bases in the U-tract sequence. Specifically, this study calculates *U*_score_ for three segments of the U-tract: the entire U-tract sequence, the first 5 bases of the U-tract, and the bases 6–8 of the U-tract. The calculation formulas are as follows [7]:

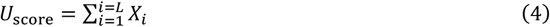

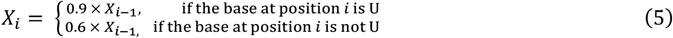 Here, *X*_0_ = 1, *i* ranges from 1 to *L*, and *L* denotes the length of the U-tract sequence.
2. The *k*-mer and C*K*SNAP features of A-tract, Hairpin, Loop, and U-tract structures. Extracted features include 1-mer, 2-mer, C1SNAP, and C2SNAP. For Loop structure within the hairpin, due to its short sequence length, only 1-mer and 2-mer features are extracted.

#### 2.1.3 Feature selection methods

Excessive features extracted from sequences may include irrelevant or redundant information, adversely affecting prediction accuracy and computational efficiency [23]. Feature selection enhances model performance by eliminating non-predictive elements and reducing runtime. This study implements a two-phase selection strategy. Phase 1 evaluates feature importance through three methods: (1) *F*-test: Measures linear correlation between features and labels via *F*-statistics and *p*-values. (2) Mutual Information (MI): Quantifies variable dependency using information entropy, where higher values indicate stronger relevance [24]. (3) Random Forest (RF): Calculates feature contributions through weighted importance scores in ensemble trees [25]. These methods represent filter-based (*F*-test/MI) and embedded (RF) approaches [26]. Phase 2 employs Incremental Feature Selection [27] to determine optimal feature quantity. Starting with the top-ranked feature, we sequentially add subsequent features, identifying the subset yielding peak regression performance. This progressive evaluation pinpoints the most effective feature combination for terminator strength prediction.

#### 2.1.4 Prediction model evaluation

In this study, 5-fold cross-validation was employed to evaluate the terminator strength prediction model. To assess model performance, two commonly used metrics for regression problems were used: Root Mean Squared Error (*RMSE*) and Coefficient of Determination (*R*^2^). *RMSE* reflects the average difference between predicted and actual values, with smaller values indicating lower error. *R*^2^ represents the proportion of variance in the data explained by the model, with values closer to 1 indicating better fit. These metrics collectively provide a comprehensive evaluation of the model’s predictive accuracy. The flowchart for the construction of terminator strength prediction model is illustrated in **Figure 1A**.

**Figure 1.**
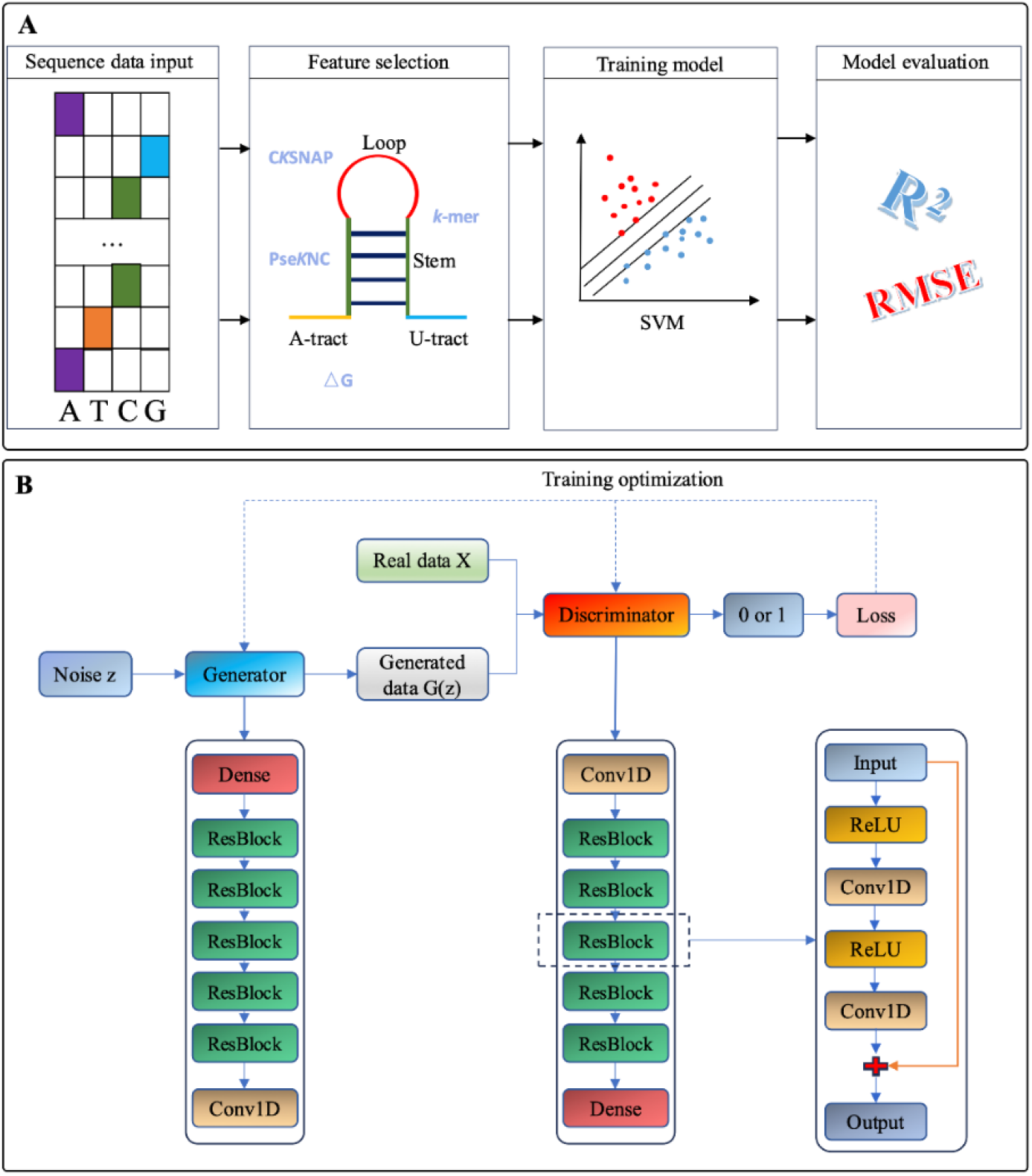
The flowchart for the constructed models. (A). Construction of terminator strength prediction model. (B). The architecture of the GAN model for terminator generation, illustrating the structures of the generator, discriminator, and Resblock.

### 2.2 Construction of generation model for terminator sequence

#### 2.2.1 Data source

Data Collection Criteria: The terminators used in this study belong to intrinsic terminators. Most intrinsic terminator sequences in the *E. coli* genome consist of three structural elements: an A-tract, a hairpin structure, and a U-tract. Therefore, the construction of the training set for intrinsic terminators followed three main constraints: (1) The terminator must be intrinsic. For synthetic terminators, only those with termination efficiency greater than 0.5 under the measurement system were included. (2) The RNA hairpin structure of the terminator must have no more than one mismatched base pair in the stem region, which must be a GU pairing. Gap pairings (unpaired regions within the stem) are not allowed. (3) The stem of the RNA hairpin structure must be ≤12 base pairs (bp) in length, and the loop must be ≤8 bp in length.

The intrinsic terminator data collected in this study were primarily sourced from the following references: (1) Intrinsic terminators with experimentally measured termination strengths from Chen et al. [6]. For synthetic terminators in this reference, only those with termination efficiency >0.5 were included. (2) Intrinsic terminator sequences from the RegulonDB database [28]. (3) Experimentally validated intrinsic terminator sequences from Dar et al. [29]. (4) Intrinsic terminator sequences from Cambray et al. [5].

After removing duplicates, the collected *E. coli* intrinsic terminators were filtered according to the three criteria above. Each terminator sequence was structured as follows: (1) The A-tract contains 8 bp. (2) The hairpin structure includes 20 bp (comprising the left half of the stem base-pairing sequence and the loop sequence; missing regions were filled with the placeholder character “P”). (3) The U-tract contains 12 bp. The final dataset comprised 641 *E. coli* intrinsic terminator sequences.

#### 2.2.2 The WGAN-GP model

Generative Adversarial Networks (GANs) were first proposed by Goodfellow et al. in 2014 [30] and have since been improved by Arjovsky et al. (Wasserstein GAN, WGAN) [31] and Gulrajani et al. (WGAN with Gradient Penalty, WGAN-GP) [32]. In this study, we adopted WGAN-GP as the core framework for terminator generation, aiming to achieve stable training while producing biologically meaningful terminator sequences.

The architecture of the WGAN-GP model, including its generator and discriminator, is illustrated in **Figure 1B**. Both components are constructed based on ResBlock modules. The generator takes a 128-dimensional random noise vector as input. This input is first processed through a Dense Layer, followed by five ResBlock modules. Finally, a series of convolutional neural networks (CNNs) with softmax as the activation function generates the terminator sequence. The discriminator receives One-hot encoded terminator sequences as input. These sequences are passed through a series of CNNs, processed by five ResBlock modules, and then fed into a dense neural network to output a judgment on the authenticity of the samples.

Within each ResBlock module, the input signal sequentially passes through two substructures (each containing a ReLU activation function and a 1D convolutional network). The output is then multiplied by a scaling factor *r* (*r* = 0.3 in this study) and added to the original input to produce the final output of the module. To enhance the discriminator’s ability to distinguish between real and generated samples, thereby more effectively updating the generator’s parameters, the discriminator was trained five times for every single training iteration of the generator.

### 2.3 Measurement of fluorescence intensity

To characterize the termination efficiency of different terminator sequences, fluorescence intensity was measured using a series of pGR plasmids (pGR1∼19) in *E. coli* MG1655. Plasmid construction was based on the template pGR0, with terminators inserted into the intergenic region between *eGFP* and *mRFP1* (**Figure S1**). The constructs were created via PCR amplification, gel extraction, and transformation into DH5α competent cells. After colony PCR and sequencing verification, the pGR plasmids were then transformed into MG1655 competent cells. Following induction of expression, the fluorescence intensity of GFP (excitation 488 nm, emission 517 nm) and RFP (excitation 560 nm, emission 650 nm) was measured. Details on the primers used, plasmid construction, and strain information can be found in **Supplementary Tables S1-S3**.

### 2.4 Calculation of termination efficiency

The transcription termination ability of *E. coli* terminators is quantified using termination efficiency (TE). In the experimental design, a dual-fluorescent reporter system was constructed using *eGFP* (green fluorescent protein gene) and *mRFP1* (red fluorescent protein gene), with the terminator insertion site located in the intergenic region between these two reporter genes (**Figure S1**). The terminational rate (*TR*) of a terminator is quantified by calculating the ratio of downstream RFP fluorescence intensity to upstream GFP fluorescence intensity. The *TR* without a terminator insertion, *TR*_ref_, serves as the reference value. The formula for calculating *TE* is as follows[5]:

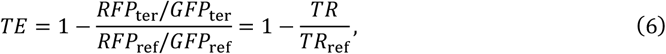

where *RFP*_ter_ and *GFP*_ter_ represent the fluorescence intensity of the downstream RFP and the upstream GFP of the terminator, respectively, and their ratio is *TR. TR*_ref_ is obtained by calculating the average of the *TR* values under two cases: the terminator is not inserted and a random sequence is inserted.

## 3. Results

### 3.1 The prediction model for terminator strength

#### 3.1.1 Feature selection on the training dataset to construct the model

To investigate the impact of terminator sequence features on terminator strength in *E. coli*, three sequence feature extraction methods were employed: *k*-mer, Pse*K*NC (Pseudo K-tuple Nucleotide Composition), and CKSNAP (Composition of *K*-spaced Nucleic Acid Pairs). Support Vector Regression (SVR) models were trained using these features, and the prediction results are summarized in **Table 2**. Key findings include: (1) for *k*-mer features, the 4-mer model achieved the best performance with an *R*^*2*^ of 0.6406; for Pse*K*NC features, Pse4NC yielded the highest *R*^*2*^ (0.6294); For C*K*SNAP features, five variants (C1SNAP to C5SNAP) were tested, achieving an *R*^*2*^ of 0.6395.

**Table 2.**
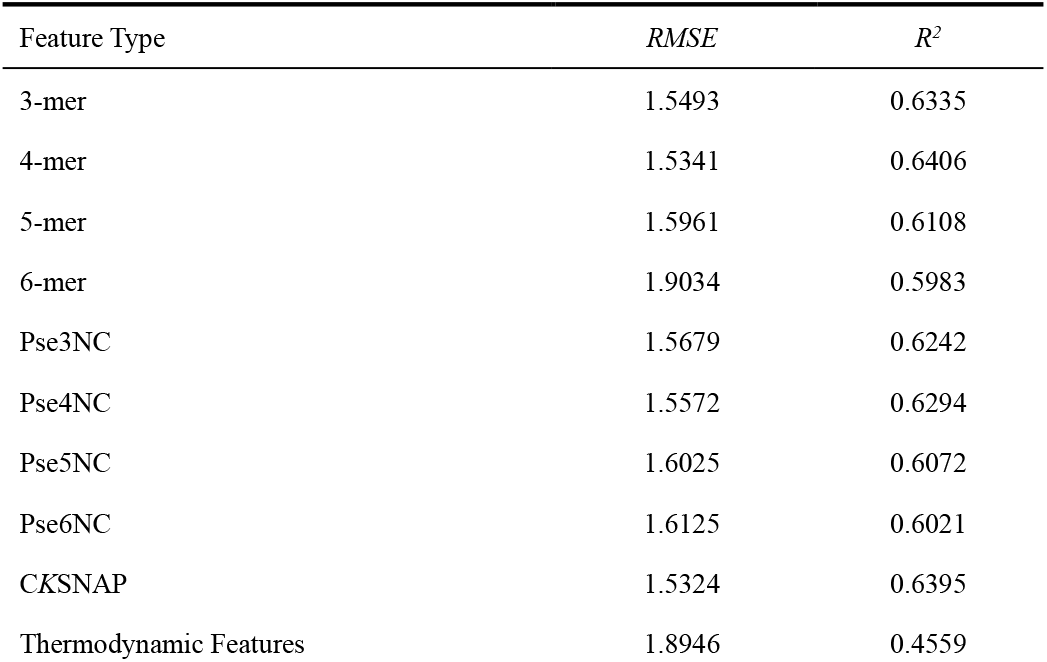
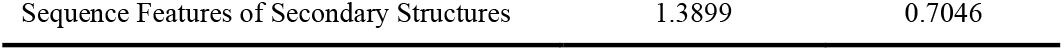
Prediction results of different features for terminators.

Additionally, thermodynamic features and RNA secondary structure sequence features were used to train SVR models. Results showed: (1) models using thermodynamic features performed poorly (*R*^*2*^ =0.4559); (2) models incorporating sequence features of RNA secondary structures achieved the highest *R*^*2*^ (0.7046), outperforming sequence-only features.

To further improve prediction accuracy, six feature sets were constructed by combining selected features (**Table 3**). Feature selection methods (*F*-test, Mutual Information, Random Forest) were applied to identify optimal sets. **Table 3** shows that Set1 (3-mer + Thermodynamic + Secondary Structure features) achieved the best performance (*R*^*2*^ = 0.7204). Thus, this combination was selected as the final model for predicting terminator strength in *E. coli*.

**Table 3.** Prediction results for six feature sets.

| Feature Set | Feature Combination | $RMSE$ | $R^2$ |
| --- | --- | --- | --- |
| Set1 | 3-mer + Thermodynamic + Secondary Structure | 1.3511 | 0.7204 |
| Set2 | 4-mer + Thermodynamic + Secondary Structure | 1.3730 | 0.7118 |
| Set3 | CKSNAP + Thermodynamic + Secondary Structure | 1.3531 | 0.7189 |
| Set4 | Pse4NC + Thermodynamic + Secondary Structure | 1.3788 | 0.7085 |
| Set5 | 3-mer + 4-mer + Thermodynamic + Secondary Structure | 1.3740 | 0.7112 |
| Set6 | 3-mer + 4-mer + CKSNAP + Thermodynamic + Secondary Structure | 1.3624 | 0.7153 |

#### 3.1.2 Comparison of prediction results with biophysical model on the test dataset

To evaluate the performance of our terminator strength prediction model, we compared it with the biophysical model proposed by Chen et al. [6] using the test dataset. The results demonstrated that our model achieved an *R*^2^ value of 0.7002 and an *RMSE* of 1.3513, closely aligning with its training set performance. In contrast, Chen’s biophysical model yielded significantly lower metrics, with an *R*^2^ of 0.2907 and an *RMSE* of 2.0786. These results confirm that our model substantially outperforms the biophysical approach in both evaluation metrics, highlighting its improved predictive accuracy for terminator strength.

### 3.2 The generation model for terminator sequence

This study employs the Wasserstein Generative Adversarial Network with Gradient Penalty **(WGAN-GP)** model to learn from intrinsic terminator sequences and generate novel terminator sequences. To evaluate the similarity between the data distribution of WGAN-GP-generated terminators and the training dataset, two methods were applied: sequence logo plots and *k*-mer sequence feature analysis.

#### 3.2.1 Evaluation based on Logo plots

Sequence logo plots were used to visualize the A-tract and U-tract structures of generated terminators and real terminators in the training set (**Figure 2**). Key observations include: (1) The 3’-end 5 bp of the A-tract in both generated and real terminators is enriched with adenine (A). (2) The 5’-end 6 bp of the U-tract in both datasets is enriched with uracil (U). (3) At each position, the nucleotide composition of generated terminators closely matches that of real terminators, demonstrating the reliability of the WGAN-GP model in mimicking natural sequence distributions.

**Figure 2.**
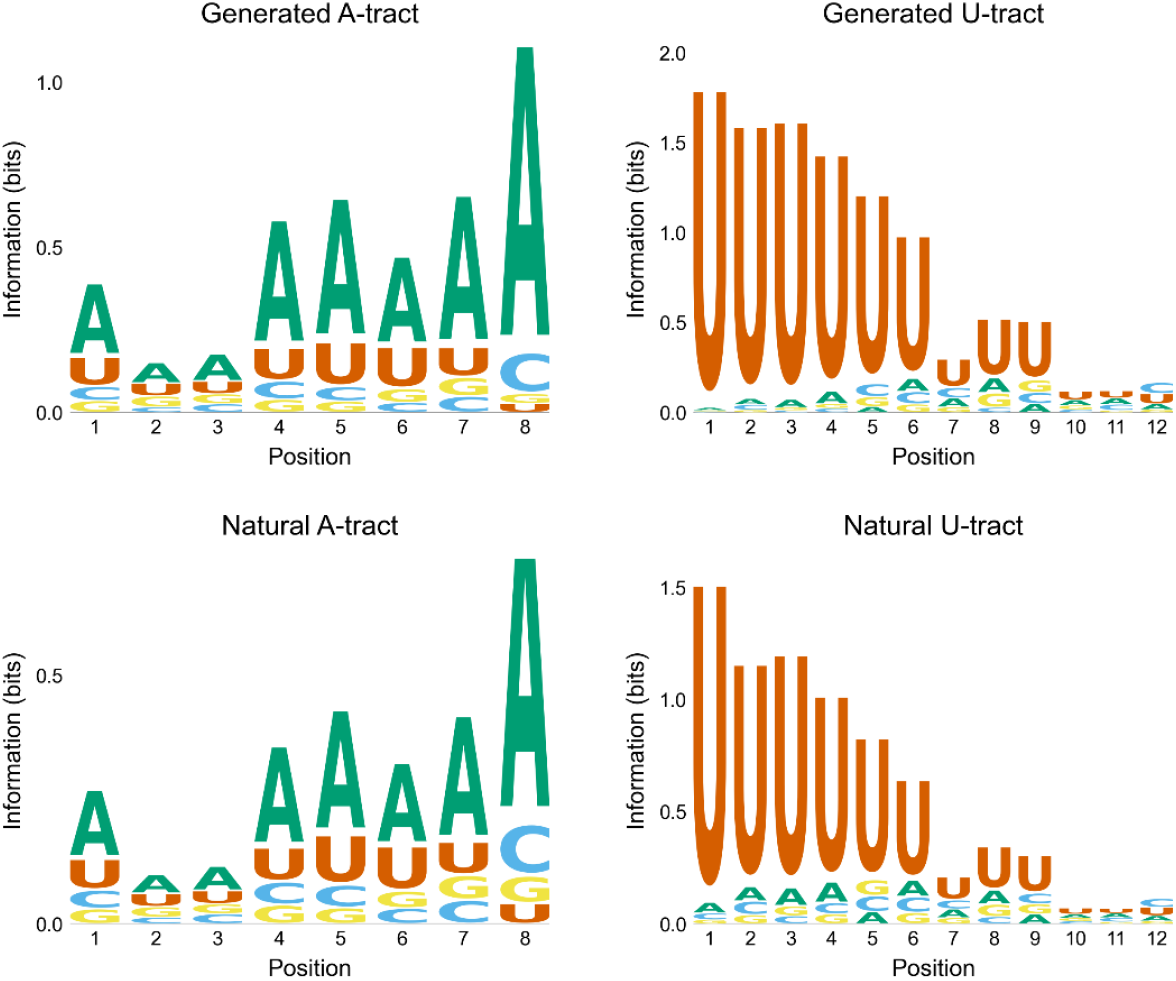
Sequence logo plots of A-tract and U-tract structures for generated terminators and real terminators in the training set.

#### 3.2.2 Evaluation based on k-mers

To assess the performance of the *E. coli* terminator generation model at the sequence feature level, *k*-mer features (2-mer, 3-mer, 4-mer, 5-mer, and 6-mer) were analyzed for the following regions: A-tract, Hairpin, U-tract, and full sequence. The Pearson correlation coefficient (*PCC*) between the *k*-mer features of generated and real terminators was calculated (**Table 4**). Results show: (1) *PCC* decreases with increasing *k*-value across all structural regions. (2) U-tract exhibits the highest correlation, with *PCC* > 0.98 for all *k*-mers, likely due to the strong U-base enrichment in its first six positions. (3) Hairpin shows the lowest correlation, but even its 6-mer *PCC* reaches 0.90. (4) These results confirm that WGAN-GP-generated terminators share high distributional similarity with real terminators.

**Table 4.**
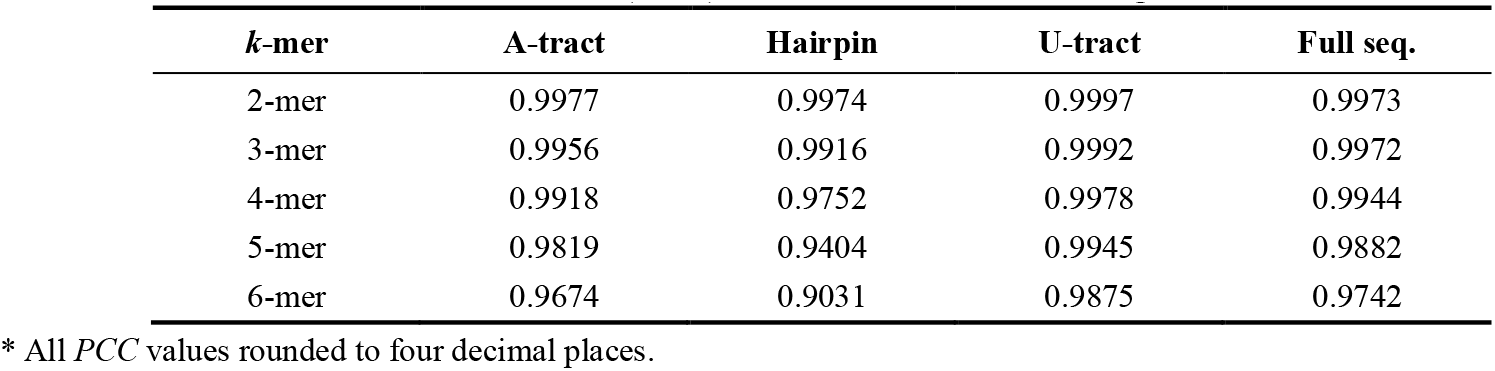
Pearson correlation coefficients (*PCC*) of *k*-mer features between generated and real terminators *.

### 3.3 Termination efficiency characterization for generated terminators

Based on the constructed *E. coli* terminator strength prediction model, this study predicted the termination strength of WGAN-GP-generated terminators and selected 20 high-efficiency candidates. Through experimental validation, 18 synthetic terminators were successfully constructed and subjected to fluorescence detection. The fluorescence intensity values of green fluorescent protein (GFP) and red fluorescent protein (RFP) for these 18 synthetic terminators, the reference terminator BBa_B0010 (a commonly used high-efficiency terminator in Synthetic Biology), **Control1** (insertion of a random sequence between *eGFP* and *mRFP1*), and **Control2** (no insertion) are shown in **Figure 3**. The results indicate that the RFP fluorescence intensity of the 18 synthetic terminators and BBa_B0010 is significantly lower than that of Control1 and Control2, demonstrating effective termination of upstream *eGFP* transcription.

**Figure 3.**
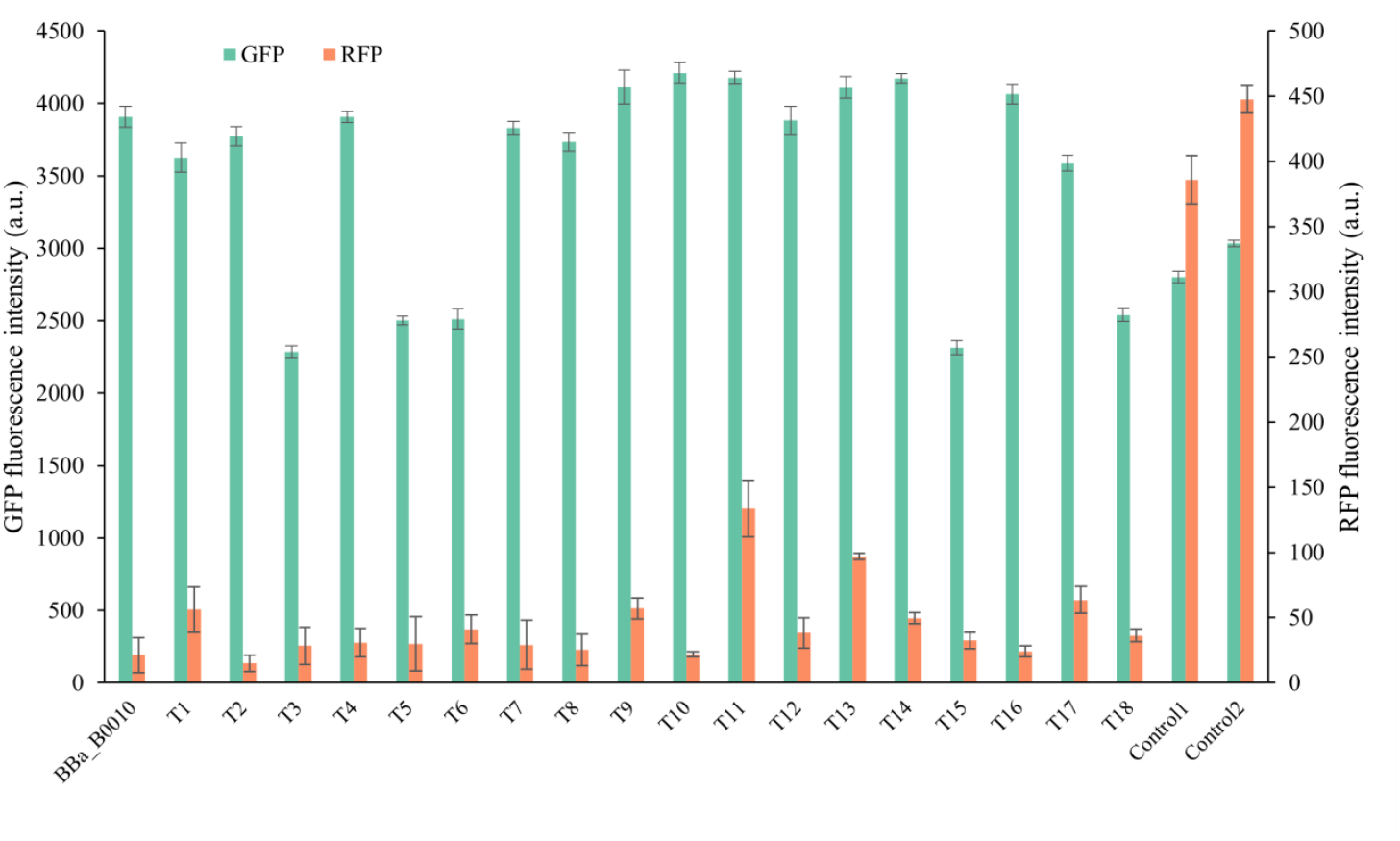
GFP and RFP fluorescence intensities of generated terminators. The GFP fluorescence intensity and the RFP fluorescence intensity were measured under different terminators. Here, Bba_B0010 represents a natural terminator, T1-T18 are the generated terminators, Control1 refers to the insertion of a random sequence between *eGFI* and *mRFP1*, and Control2 indicates no insertion between the two. A lower ratio of RFP fluorescence intensity to GFP fluorescence intensity indicates higher termination efficiency of the terminator.

To quantify termination efficiency, the termination rates of Control1 and Control2 (representing scenarios without terminators) were averaged as the reference baseline. Termination efficiency values for BBa_B0010 and the 18 synthetic terminators are listed in **Table 5**. Key findings include: (1) 13 out of 18 terminators (72%) achieved termination efficiency > 0.90. (2) 16 out of 18 terminators (89%) achieved termination efficiency > 0.85. (3) 5 synthetic terminators (T2, T7, T8, T10, T16) exhibited efficiency comparable to BBa_B0010 (0.96). These results validate the potential of the WGAN-GP model to explore the sequence space of *E. coli* terminators and confirm the accuracy of the terminator strength prediction model in screening high-efficiency candidates.

**Table 5.**
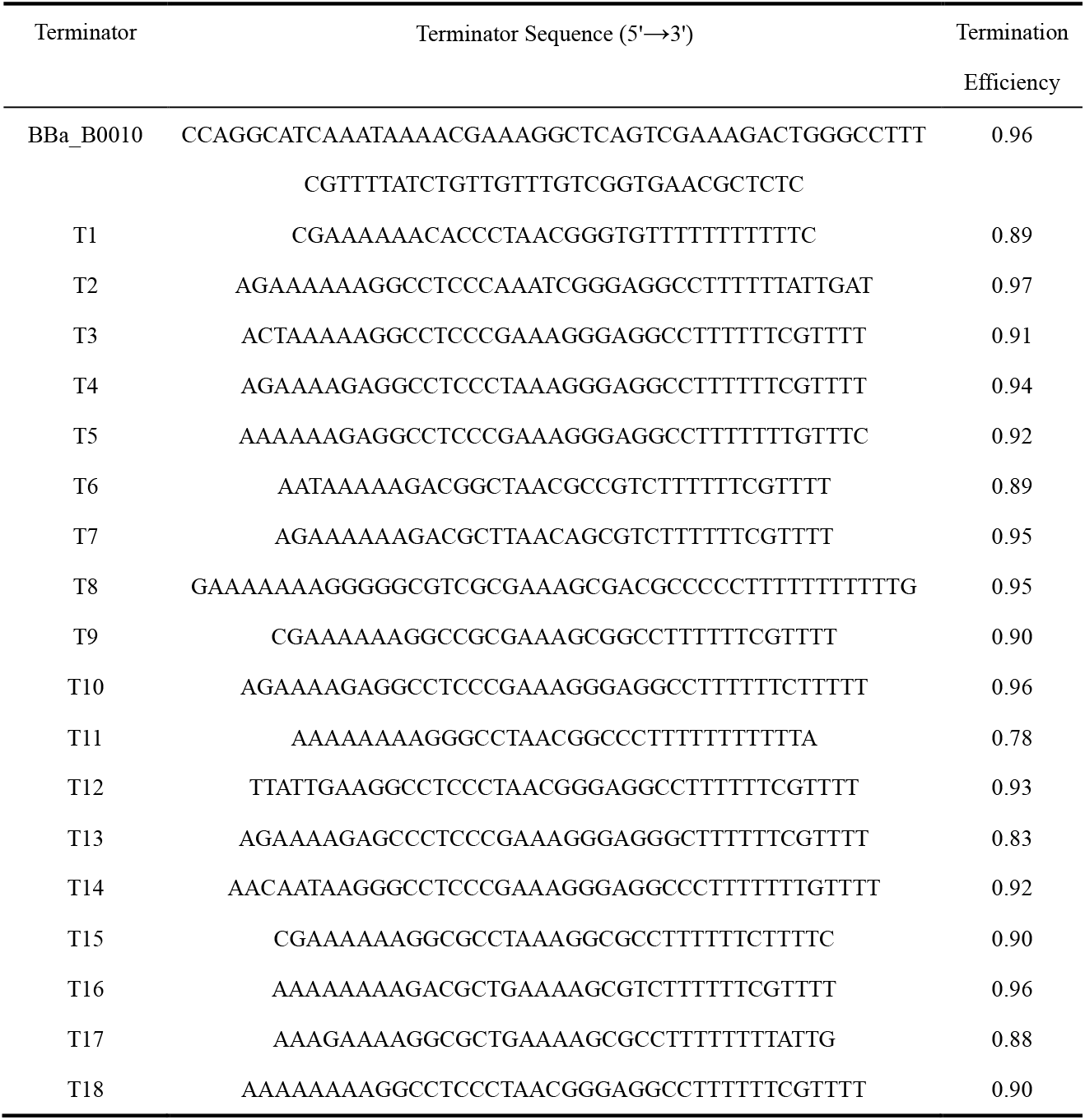
Termination efficiency of generated terminators.

## 4. Discussion

The development of computational tools for predicting and designing genetic regulatory elements has become a cornerstone of synthetic biology, yet the characterization of terminators has lagged behind that of promoters and ribosome-binding sites (RBS). This study addresses this gap by integrating machine learning and generative adversarial networks (GANs) to predict terminator strength and generate novel high-efficiency terminators for *Escherichia coli*. Our approach demonstrates the feasibility of leveraging computational models to streamline the design of functional genetic parts, reducing reliance on labor-intensive experimental screening. Below, we contextualize our findings within the broader field, highlight methodological innovations, and discuss implications for future research.

The support vector regression (SVR) model developed here, which combines 3-mer sequence features, thermodynamic properties, and RNA secondary structure features, achieved a predictive performance (*R*^*2*^ = 0.7204) superior to models relying solely on sequence or thermodynamic features. This aligns with prior studies emphasizing the importance of integrating multiple feature types for accurate prediction of transcriptional termination. For instance, it was demonstrated that RNA secondary structure stability, particularly in hairpin loops, significantly influences terminator efficiency [33], a finding corroborated by our observation that secondary structure-derived features contributed significantly to model accuracy. Similarly, the inclusion of thermodynamic parameters such as Δ*G* (minimum free energy) echoes work by Cui et al. [34], who highlighted free energy as a critical determinant of termination efficiency (TE). Recent work by Zhai et al. [7] further validated that terminators with longer stems and perfect U-tracts exhibit significantly higher TE, underscoring the necessity of structural stability in termination efficiency—a principle mirrored in our feature extraction strategy.

The application of Wasserstein GANs with gradient penalty (WGAN-GP) to generate synthetic terminators represents a novel contribution to the field. While GANs have been used to design promoters [8] and RBS sequences [35], their use for terminator generation remains underexplored. Our evaluation via sequence logos and *k*-mer Pearson correlation coefficients (*PCC* > 0.90 for all features) confirmed that WGAN-GP-generated terminators closely mimic the sequence distribution of natural terminators, particularly in conserved regions like U-tracts. This mirrors findings from DNA generative models, such as those by Killoran et al. [36], where GANs successfully captured important features in DNA sequences, such as exon splice site signals and nucleotide complementarity. Notably, the high *PCC* values for U-tract *k*-mers (*PCC* > 0.98) suggest that the model effectively learned the strong bias toward uracil enrichment in this region—a hallmark of bacterial intrinsic terminators. However, the lower correlation in hairpin structures (*PCC* = 0.90 for 6-mer) underscores the challenge of modeling complex RNA folding patterns, a limitation also observed in RNA structure prediction tools like RNAfold [37]. Experimental validation of the 18 generated terminators revealed that 72% exhibited termination efficiency > 0.90, with five matching the performance of the canonical high-efficiency terminator BBa_B0010. These results underscore the practical utility of combining predictive models with generative algorithms to prioritize candidates for experimental testing. Notably, the success rate of our computational pipeline (89% of terminators exceeding 85% efficiency) compares favorably to traditional screening methods [5]. This efficiency gain is critical for scaling synthetic biology applications, such as metabolic engineering and genetic circuit design, where large libraries of terminators are required to fine-tune gene expression.

While our work advances terminator design, several limitations warrant attention. First, the model’s reliance on *E. coli* data limits its generalizability to other species, as terminator mechanisms vary across bacteria and eukaryotes [4, 38]. Expanding training datasets to include diverse organisms could enhance versatility. Second, the SVR model’s dependence on handcrafted features (e.g., *k*-mers) may overlook higher-order sequence interactions. Emerging deep learning architectures, such as convolutional neural networks (CNNs) or transformers, could automatically extract such patterns, as demonstrated in protein design [39, 40]. Finally, while WGAN-GP generated plausible terminators, the biological plausibility of all candidates, such as compatibility with host transcription machinery, remains to be fully explored. Integrating evolutionary constraints or energy-based refinement steps, as proposed by Sinai et al. [41], could further improve generative accuracy. Du et al. [42] demonstrated that embedding biological knowledge (e.g., motif conservation) into generative models enhances functional sequence output—a strategy that could be adapted for terminator design.

This study establishes a robust framework for computational terminator design, combining predictive modeling with generative AI to accelerate the discovery of high-efficiency terminators. By demonstrating the synergy between sequence-structure features and deep generative models, our work paves the way for similar approaches in designing other regulatory elements. Future efforts should focus on broadening taxonomic applicability, leveraging advanced neural architectures, and integrating automated experimental validation platforms to fully realize the potential of computational synthetic biology.

## Supporting information

This article contains supporting information.

## Acknowledgements

This work was supported by the grant from the National Natural Science Foundation of China (31971184).

## Supporting information

**Figure S1.**
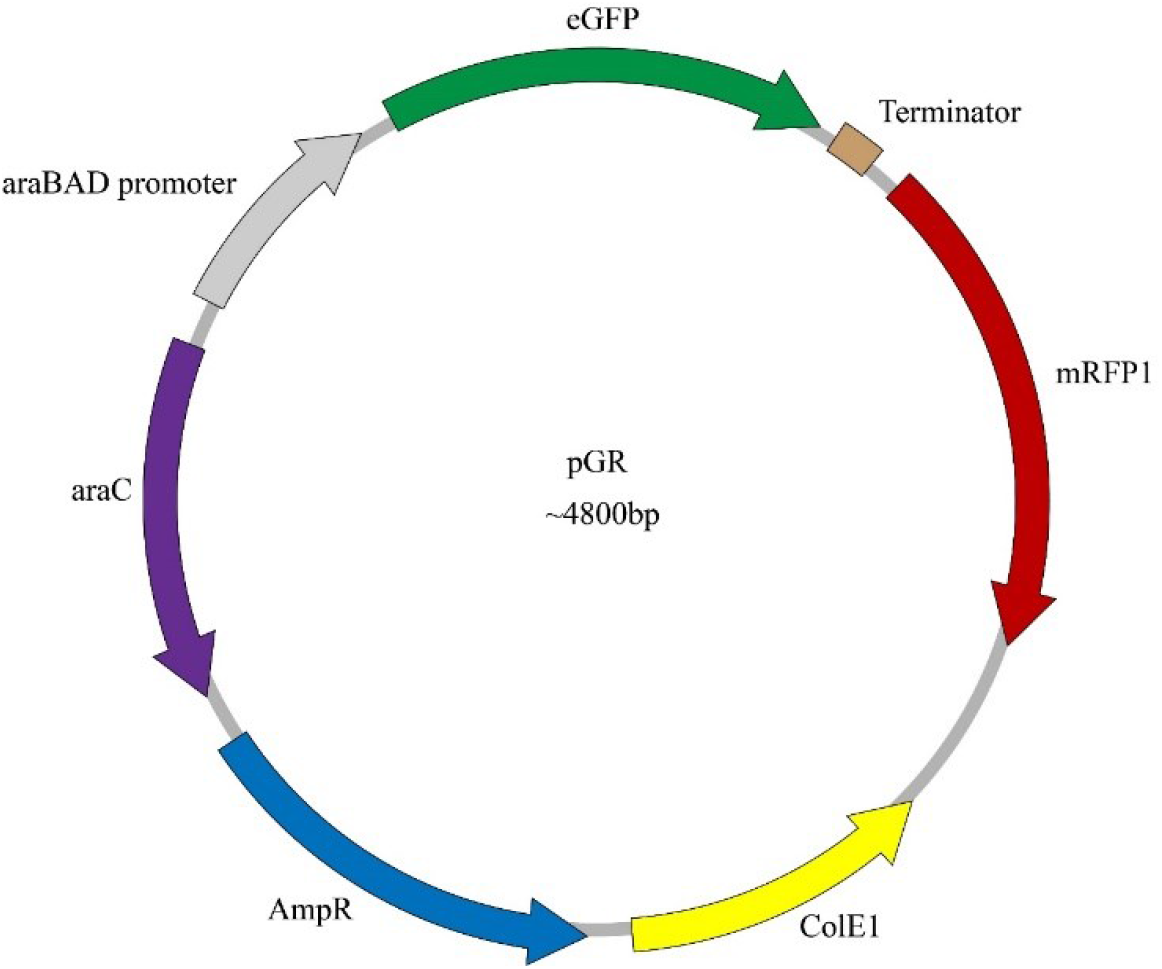
Plasmid map of pGR series.

**Table S1.**
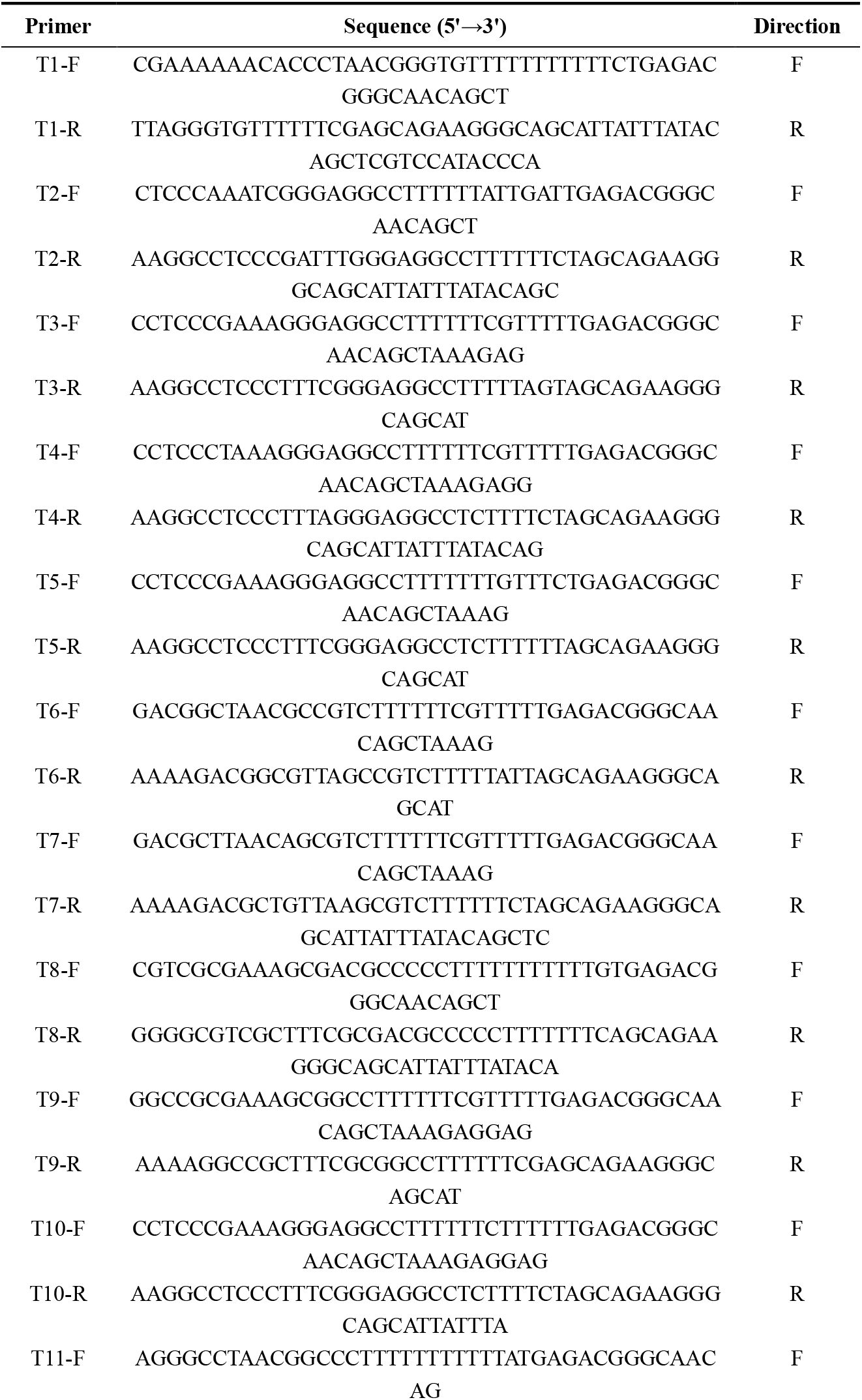
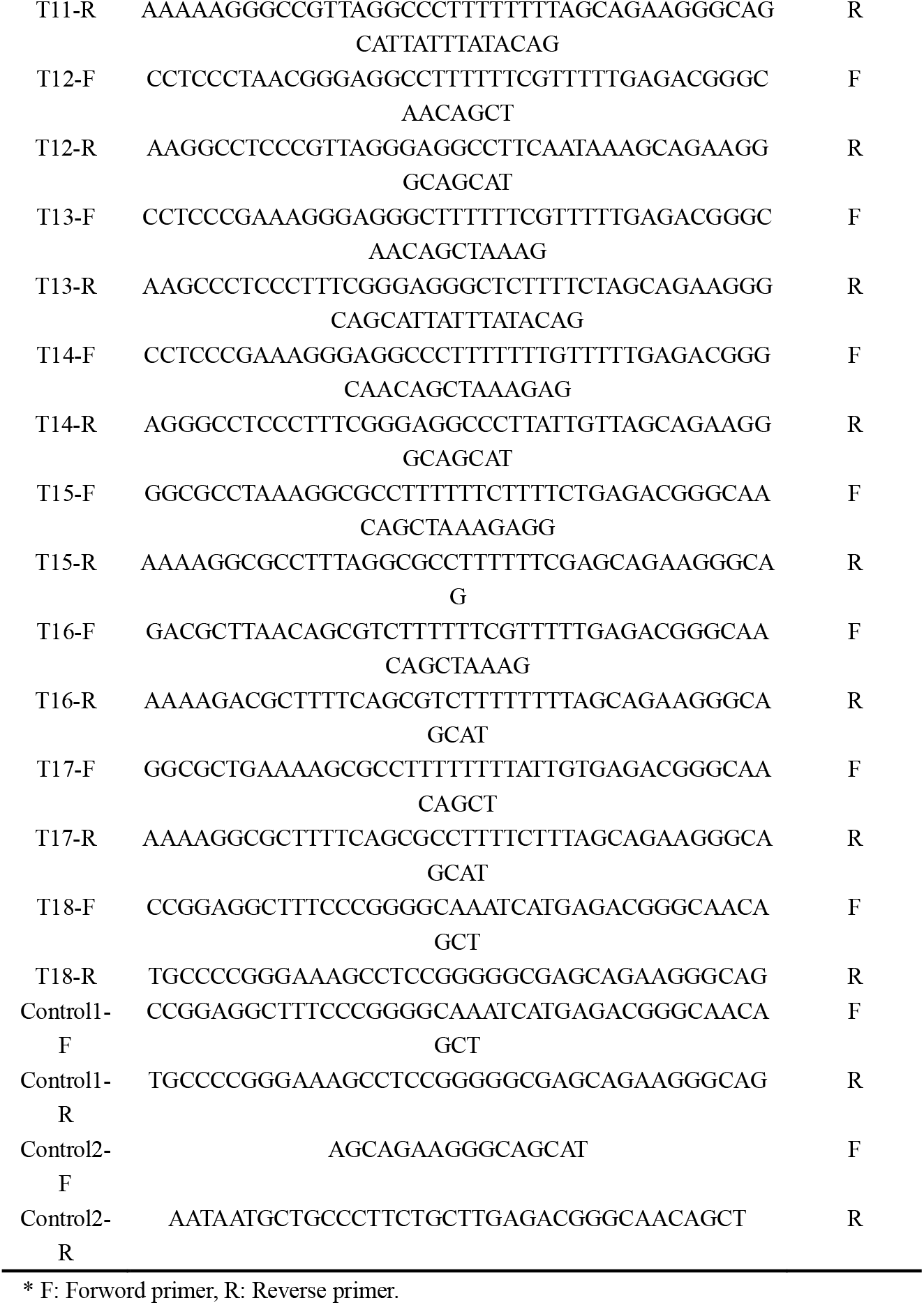
The primers used in this study*.

**Table S2.**
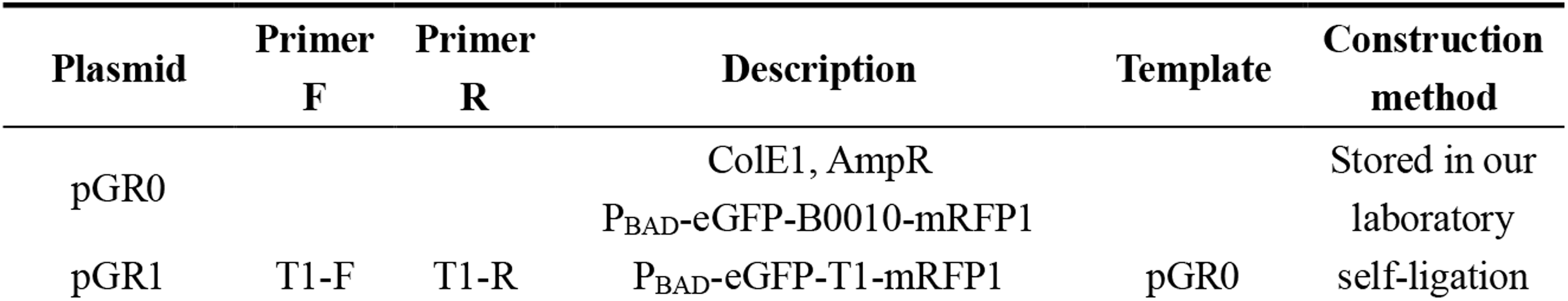
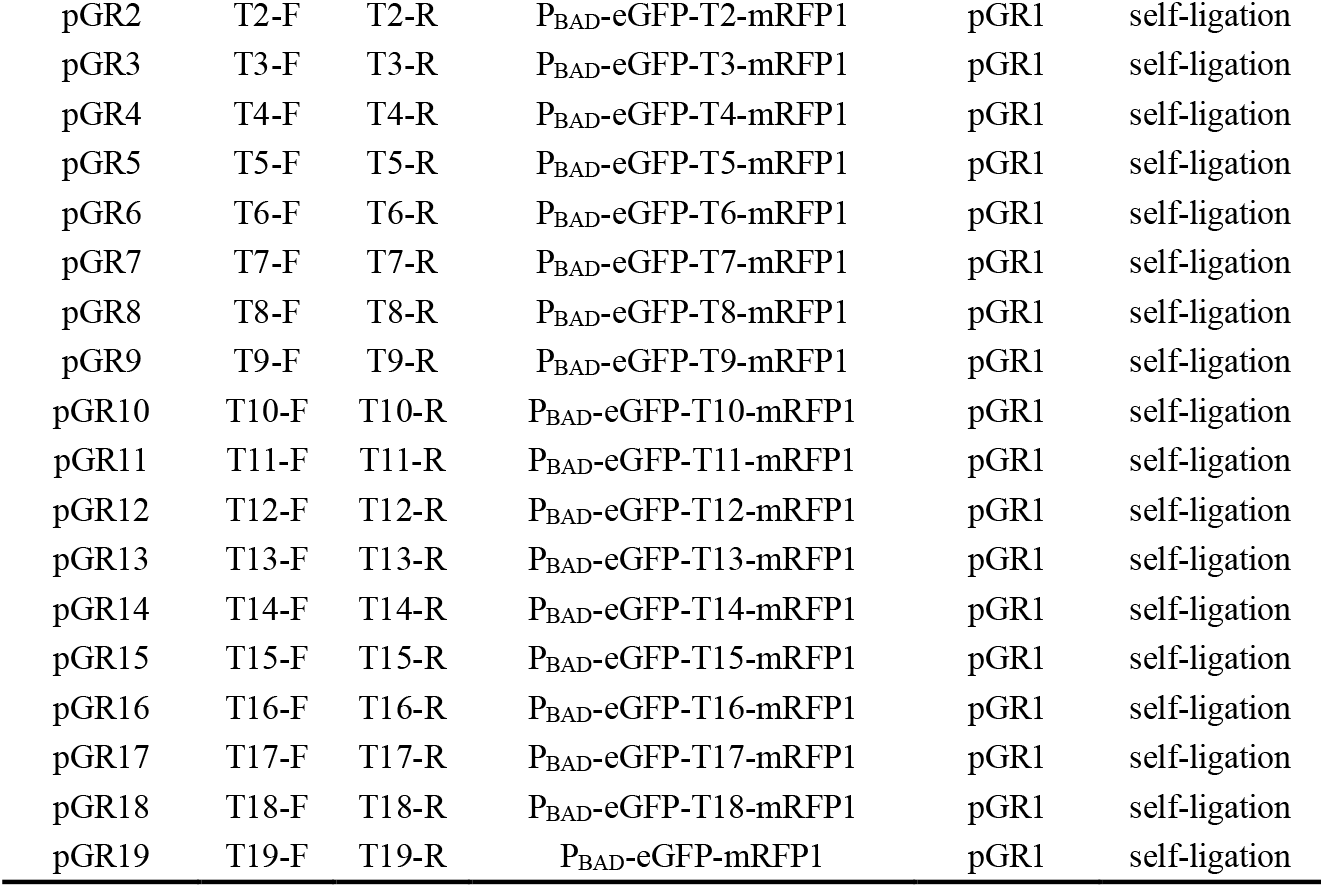
Plasmid construction details in this study.

**Table S3.** The *E. coli* strains used in this study.

| Strain | Genotype | Source |
| --- | --- | --- |
| MG1655 | Standard strain *E. coli* MG1655 | Stored in our laboratory |
| DH5α | Standard strain *E. coli* DH5α | Stored in our laboratory |
| LJ0 | MG1655/ pGR0 | Constructed in this study |
| LJ1 | MG1655/ pGR1 | Constructed in this study |
| LJ2 | MG1655/ pGR2 | Constructed in this study |
| LJ3 | MG1655/ pGR3 | Constructed in this study |
| LJ4 | MG1655/ pGR4 | Constructed in this study |
| LJ5 | MG1655/ pGR5 | Constructed in this study |
| LJ6 | MG1655/ pGR6 | Constructed in this study |
| LJ7 | MG1655/ pGR7 | Constructed in this study |
| LJ8 | MG1655/ pGR8 | Constructed in this study |
| LJ9 | MG1655/ pGR9 | Constructed in this study |
| LJ10 | MG1655/ pGR10 | Constructed in this study |
| LJ11 | MG1655/ pGR11 | Constructed in this study |
| LJ12 | MG1655/ pGR12 | Constructed in this study |
| LJ13 | MG1655/ pGR13 | Constructed in this study |
| LJ14 | MG1655/ pGR14 | Constructed in this study |
| LJ15 | MG1655/ pGR15 | Constructed in this study |
| LJ16 | MG1655/ pGR16 | Constructed in this study |
| LJ17 | MG1655/ pGR17 | Constructed in this study |
| LJ18 | MG1655/ pGR18 | Constructed in this study |
| LJ19 | MG1655/ pGR19 | Constructed in this study |

